# AdiY Acts as a Cytoplasmic pH-Sensor via Histidine Protonation to Regulate Acid Stress Adaptation in *Escherichia coli*

**DOI:** 10.1101/2025.10.03.680266

**Authors:** Giovanni Gallo, Sophie Brameyer, Sonja Kuppermann, Sabine Schneider, Pavel Kielkowski, Kirsten Jung

## Abstract

The arginine-dependent acid resistance (Adi) system is a vital component that enables *Escherichia coli* and other enterobacteria to withstand the extreme acidity in the human gastrointestinal tract. It consists of the proton-consuming decarboxylation of arginine, catalyzed by AdiA, and the uptake of arginine, as well as the excretion of the more alkaline agmatine, catalyzed by the antiporter AdiC. The corresponding genes *adiA* and *adiC* are induced in *E. coli* under acidic conditions (pH < 5.5), a process that is tightly regulated by the AraC/XylS transcriptional activator AdiY. Here, we show that the pH-sensing mechanism of AdiY functions through the protonation of two histidines (His34 and His60) in the N-terminal domain. Replacing these histidine residues with alanine, glutamine or aspartate abolishes the pH-dependent activation of AdiY, both *in vivo*, as demonstrated by promoter-reporter assays, and *in vitro*, as indicated by the loss of DNA-binding activity detected by surface plasmon resonance spectroscopy. Biochemical analyses of purified wild-type AdiY using size-exclusion chromatography and intrinsic tryptophan fluorescence revealed a pronounced and reversible pH-dependent conformational change that does not occur in the pH-sensing–deficient AdiY variant. A model is proposed in which AdiY forms a monomer at physiological pH. At a lower intracellular pH, the protonation of histidine in AdiY causes a conformational change that leads to the binding of AdiY as a tetramer to the DNA. This work elucidates the molecular mechanism of a one-component signal transduction system that combines both sensory and responsive functions.

**Importance:** Throughout their life, *Escherichia coli* and other bacteria may encounter acidic environments, for example, when passing through the human stomach. Their chances of survival under these conditions depend on the number and efficiency of acid resistance systems. Although many acid resistance mechanisms have been extensively studied, the molecular mechanism by which bacteria sense low pH is not yet fully understood. This study demonstrates that the transcription factor AdiY acts as a direct pH sensor by using two histidines to detect intracellular acidification in *E. coli*. When these histidines become protonated, AdiY changes its conformation and activates genes that support cell survival under acid stress. These findings not only reveal a new way in which bacteria can perceive extremely low pH environments but also provide the basis for the development of AdiY as a pH reporter.

## Introduction

Microorganisms must constantly monitor changes in the proton concentration (pH) in their environment and respond accordingly (1). An increase in proton concentration (low pH) can lead to acid stress, which causes proteins, nucleic acids, and membranes to become protonated, disrupting their structure and function. Bacteria such as *Escherichia coli* that are neutralophilic face extreme acid challenges during passage through the stomach and gut of a host, where hydrochloric acid and fermentation products, such as short-chain fatty acids, lower both the extracellular and intracellular pH of the bacteria (2, 3). To survive these conditions, *E. coli* and other enteric bacteria have evolved robust acid resistance (AR) systems (4, 5). Acquisition of AR is indeed an essential trait to survive acidic environments; for example, survival in the gastrointestinal tract requires the ability to buffer or expel excess protons. These adaptive responses allow bacteria to maintain a viable cytoplasmic pH range despite extreme external acidity (6).

*E. coli* employs multiple acid resistance (AR) systems, with the most effective enzyme-based mechanisms being the glutamate-dependent (AR2/Gad), arginine-dependent (AR3/Adi), and lysine-dependent (AR4/Cad) systems. In each system, a cytoplasmic decarboxylase consumes a proton while converting its substrate (glutamate, arginine or lysine, respectively) to the corresponding amine (γ-aminobutyrate, agmatine or cadaverine, respectively), and a specific antiporter (GadC, AdiC, CadB) exports the amine in exchange for its cognate amino acid (4). This coupled decarboxylation– antiport cycle reduces the intracellular proton load, thereby increasing the cytoplasmic pH and, in the case of the Cad and Adi systems, also elevating the external pH by the secretion of the alkaline cadaverine and agmatine, respectively (7–9). A fourth acid resistance system (AR1) operates via the Sigma factor S (RpoS) and indirectly affects the F_1_F_o_-ATPase proton pump, and AR5, the ornithine decarboxylase system, plays only a minor role in laboratory strains (4, 10). Each AR system is induced under specific conditions and pH ranges, and an intricate network of sensory and regulatory proteins regulates its expression. The Gad (AR2) system, for example, is activated by a cascade of sensors and transcription factors and is essential for survival at pH levels of 2–3. In all cases, acid resistance predominantly relies on the proton-consuming reactions to restore intracellular pH homeostasis (11, 12). The regulation of AR systems often involves transcription factors from the AraC/XylS family (13). For example, the genes of the Gad system are controlled by the AraC-like regulators GadE, GadW and GadX (14). Transcription factors of the same family, such as AppY and EnvY, modulate gene expression under anaerobic or starvation conditions (15, 16). AdiY, which is in the focus of this study, is the activator of the Adi AR system and also a member of the AraC/XylS family (13, 17–21).

AdiY is a transcriptional activator that consists of 253 amino acids and binds upstream of *adiA and adiC*. It induces their expression under low pH and anaerobic conditions, such as those found in the stomach and gut (19). The *adiY* gene is located immediately downstream of *adiA*, and *adiA* and *adiC* are maximally induced at an external pH of 4.4 when *E. coli* is grown in an amino acid-rich medium. The Adi system is heterogeneously activated with only a fraction of the population expressing high levels of Adi components under acid stress (22). This pattern can be shifted to a more uniform expression when the copy number of AdiY is increased (22).

Most bacterial acid stress sensors are membrane-integrated proteins with an outwardly exposed sensor domain. A classic example is CadC, the activator of the Cad system. CadC has a C-terminal periplasmic domain that functions as an external pH sensor. At low external pH (pH 5.8) and in the presence of lysine, conformational changes in CadC trigger transcription of the *cadBA* operon, thereby coupling the external acid stress to gene activation (23–26).

In contrast, cytosolic regulators such as AdiY lack a membrane-linked sensor domain. This raises the question of which molecular features enable a cytosolic activator to respond to acid stress.

Bacteria, in general, are able to maintain a relatively constant internal pH (pH) when grown in a wide range of media with different external pH values [reviewed in (5, 27, 28)]. The cytoplasmic pH of *E. coli*, for example, only changes from 7.2 to 7.8 over an environmental pH range of 5.5 to 9 (29). Although the cytoplasmic membrane is impermeable to protons, some protons enter the cytoplasm through protein channels, and via transient water channels or damaged membranes (4, 30). For example, the pH of the cytoplasm drops transiently to pH 6.2 after exposure of *E. coli* to pH 5.8. However, intracellular pH rapidly returns to neutral (< 4 min) due to the intrinsic buffering capacity of the cytoplasm or alterations in the flux of other ions (27). At more severe acid stress (pH 4.4), the intracellular pH decreases by approximately one pH unit and remains unchanged. Genes coding for the Adi system are induced. The correlation between a decrease in intracellular pH and the activation of the Adi system was previously indirectly demonstrated, as the *adiA* promoter was strongly activated in a mutant lacking the AR4/Cad system or less activated after overproduction of the AR4/Cad system (22).

It is hypothesized that AdiY responds to the acidification of the cytoplasm. In this study, we combined genetic, biochemical, and structural analyses to find that the pH-dependent protonation of two histidines triggers a conformational change in AdiY, enabling its DNA binding and transcriptional activation. These studies offer new functional insights into a member of the AraC/XylS family transcription regulator, which may also be representative of other members, particularly those involved in virulence (13).

## Results

### AdiY is essential for pH-dependent activation of the AR3/Adi system in E. coli

To characterize the regulatory logic underlying the activation of the Adi system, we quantified the promoter activity of the *adiA, adiC*, and *adiY* genes using luciferase-based reporter plasmids (Figure 1A). These constructs were introduced into wild-type and Δ*adiY E. coli* MG1655 strains, and promoter activities were monitored across a range of pH values under both aerobic and microaerobic conditions (Figures 1B-D). The results showed that in the absence of AdiY, the *adiA, adiC*, and *adiY* promoters remained inactive under all tested conditions, regardless of pH or oxygen availability (Figures 1B-D). In contrast, wild-type cells showed strong induction of both the *adiA* and *adiC* promoters under acidic conditions, with activation thresholds below pH 5.8 in microaerobic environments and maximal expression at pH 4.4. Under aerobic conditions, activation of both promoters was diminished and occurred only at external pH values below 4.8 (Figures 1B and 1C). Notably, the activation profiles of the *adiA* and *adiC* promoters were nearly identical; however, the *adiA* promoter exhibited an eightfold higher level of induction across all tested conditions. In contrast, the activity of the *adiY* promoter remained consistently low under all tested conditions, with only a modest increase under microaerobic, acidic conditions (Figure 1D).

**Figure 1.**
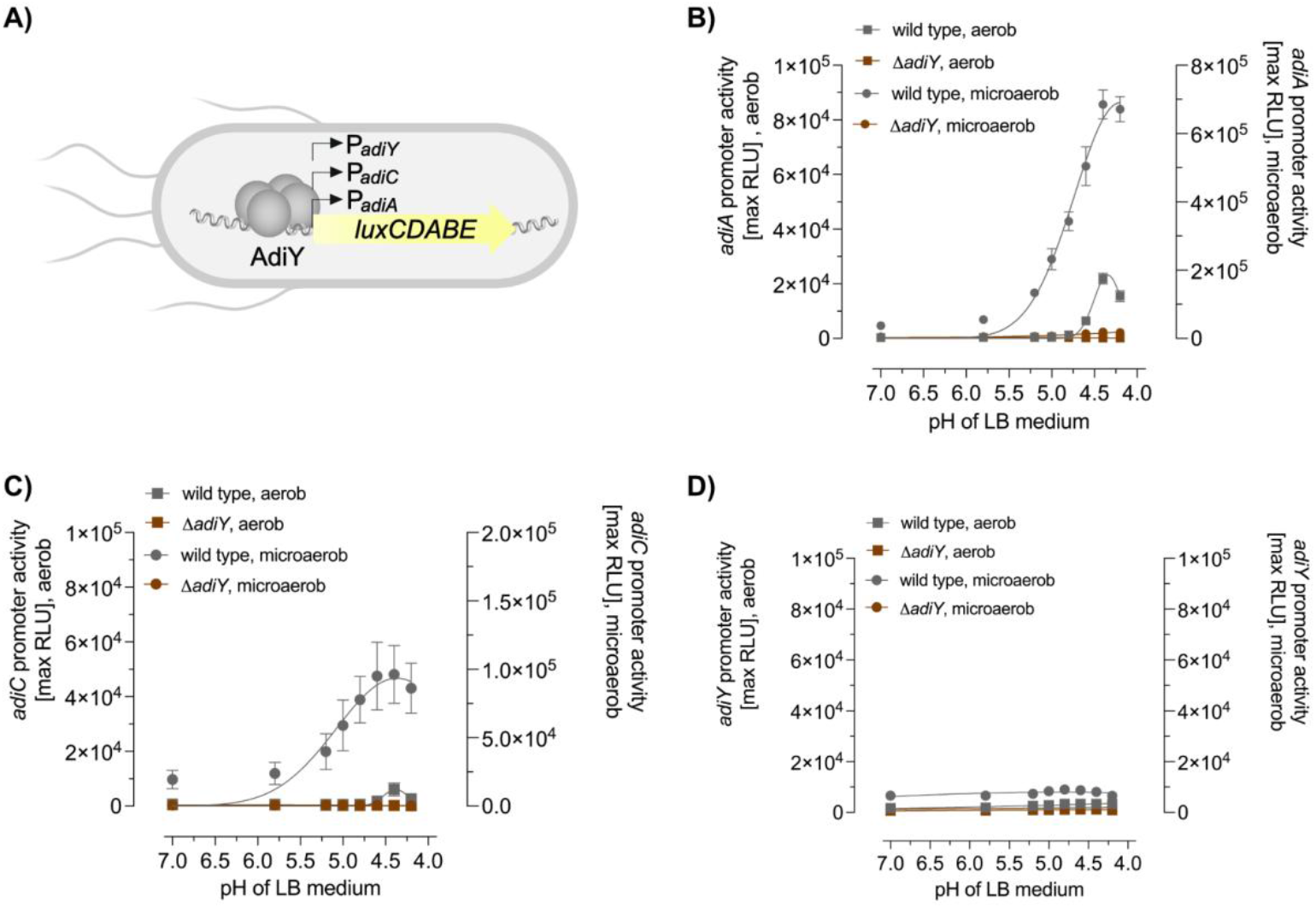
pH-dependent activation of the *adiA, adiC*, and *adiY* promoters in *E. coli* MG1655 wild-type and the *adiY* mutant. **(A)** Schematic overview of the luminescence-based reporter, in which the expression of luciferase is controlled by the *adiA* (P_*adiA*_), *adiC* (P_*adiC*_), or *adiY* (P_*adiY*_) promoter. *E. coli* MG1655 wild type and the *adiY* mutant were each transformed with the reporter plasmid pBBR1-MCS5-P_*adiA*_-*lux* **(B)**, pBBR1-MCS5-P_*adiC*_-*lux* **(C)**, and pBBR1-MCS5-P_*adiY*_-*lux* **(D)**, and were grown in LB medium adjusted to different pH values ranging from 4.0 to 7.0. Data are reported as relative light units (RLUs) in counts per second per milliliter per OD_600_, and the maximal RLU after 2 h of growth is shown. All experiments were performed three times (n=3), and error bars represent the standard deviation of the means.

These results demonstrate that activation of the *adiA* and *adiC* promoters is strictly pH-dependent and modulated by oxygen availability, with robust induction of *adiA* and *adiC* occurring under acidic, microaerobic conditions, reduced activation under aerobic conditions, and no activation in the absence of AdiY. The *adiY* promoter itself is only weakly responsive, which is consistent with its role as a constitutive or weakly inducible regulator within the system.

### Two histidine residues are essential for AdiY activation in vivo

To elucidate the molecular mechanism of AdiY-mediated pH sensing, a systematic substitution mutagenesis approach was employed to investigate the role of histidine residues in the N-terminal domain of AdiY. Histidine residues are known to change their protonation state in response to pH fluctuations, which can influence protein conformation and function (31). Each of the four histidine residues in the N-terminal domain was individually replaced with alanine, and the resulting AdiY variants were analyzed for their ability to activate the *adiA* promoter in vivo under acidic conditions, using the *E. coli* MG1655 Δ*adiY* background for comparison to wild-type AdiY expressed from a pBAD24 plasmid. Only wild-type AdiY complemented the *adiY* deletion strain and restored full activation of the *adiA* promoter at pH 4.4 (Figure 2A). In contrast, the histidine-to-alanine substitutions at positions 34 and 60 resulted in a significant loss of function, substantially impairing AdiY’s ability to respond to low pH and activate the *adiA* promoter. Additionally, the double histidine variant AdiY-H34A/H60A displayed a synergistic loss of function, retaining only ∼5% of the low pH-responsiveness observed with wild-type AdiY (Figure 2A). Similarly, a loss of function occurred when these two positions were exchanged to glutamine or aspartate (Figure S1). All variants and wild-type AdiY were produced upon induction with L-arabinose. Furthermore, neither these variants nor wild-type AdiY showed activation of the *adiA* promoter at physiological pH (pH 7.0) (Figure 2A and S1). This finding suggests that the histidine residues at positions 34 and 60 are critical for the ability of AdiY to activate the *adiA* promoter in response to acidic pH.

**Figure 2.**
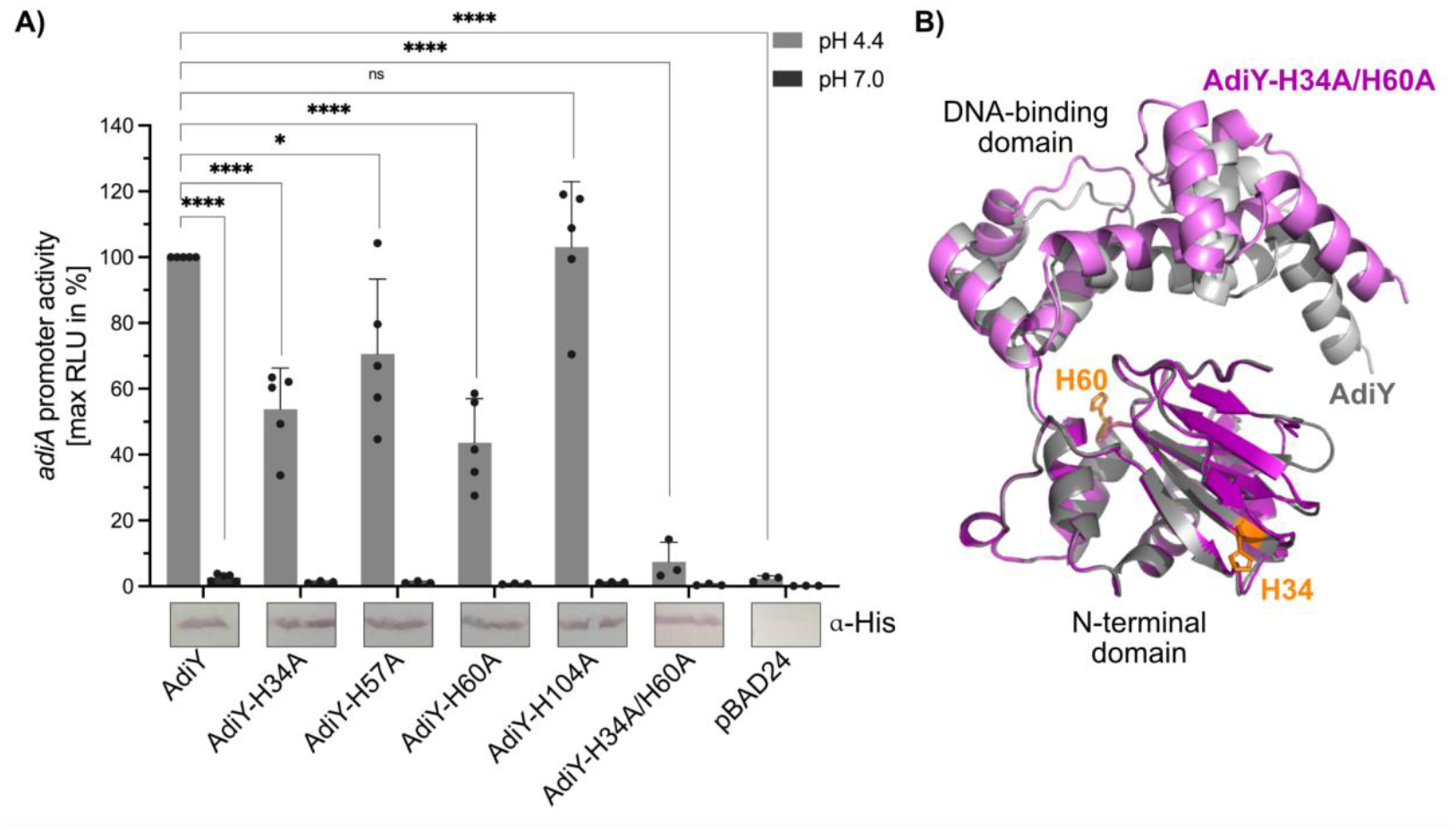
Two histidine residues are essential for AdiY-mediated activation of the *adiA* promoter under acidic conditions. **(A)** *E. coli* MG1655 Δ*adiY* was co-transformed with the reporter plasmid pBBR1-MCS5-P_*adiA*_-*lux* and plasmid-based AdiY wild-type or variants. Cells were grown in a citrate-buffered medium adjusted to pH 4.4 (grey) or pH 7.0 (dark grey) and supplemented with 0.1% L-arabinose. Data are reported as relative light units (RLUs) in counts per second per milliliter per OD_600_, and the maximal RLU after 2 h of growth is shown. All experiments were performed at least three times (n≥3), and error bars represent the standard deviation of the means. Production of AdiY wild-type or AdiY variants was confirmed by western blot analysis using antibodies against the His-tag. Protein bands correspond to a 29-kDa protein. The blot is shown split because the samples were initially loaded onto the gel in a different order. The bands were cut and rearranged to achieve the desired order in the graph. Statistics: Student’s unpaired two-sided t test; ns, not significant; ^*^ p = 0.0203; ^****^ p < 0.0001. **(B)** AlphaFold 3 structural model of the monomer of AdiY wild-type (grey) and AdiY-H34A/H60A variant (pink). Key histidine residues at positions 34 and 60 are highlighted in orange in the AdiY wild-type structural model. The figure was prepared with PyMOL (v3.1.4.1).

To gain structural insight, AlphaFold3 models were generated for both wild-type AdiY and the double histidine variant AdiY-H34A/H60A (Figure 2B). The predicted monomeric structure of wild-type AdiY (Figure 2B, grey) displays the expected AraC-family fold. His60, one of the essential histidine residues (highlighted in orange), is strategically positioned at the interface between the N-terminal regulatory and C-terminal DNA-binding domain. By contrast, the AdiY-H34A/H60A model (Figure 2B, pink) shows a different arrangement of the domains, with the N-terminal domain slightly spread out relative to the C-terminal DNA-binding domain around the H60 hinge. This structural difference suggests that these histidine residues are essential for maintaining the interdomain architecture necessary for effective transcriptional activation. Depending on the protonation state, His60 might form hydrogen bonds to Glu41 and the carbonyl oxygen of Met142. His34 is located at a β-turn in the N-terminal domain, where its protonation state also likely impacts on the structure and dynamics of its environment (Figure S6).

Collectively, these data demonstrate that two histidine residues in AdiY function as pH-sensitive molecular switches. Their protonation likely induces conformational changes that facilitate DNA-binding and enable transcriptional activation. Substitution of these residues abolishes AdiY activity *in vivo*, highlighting their critical role in acid-responsive gene regulation.

### pH-Dependent Conformational Dynamics of AdiY in vitro

To corroborate the *in vivo* finding that only wild-type AdiY, but not the AdiY-H34A/H60A mutant, can activate the *adiA* promoter in a pH-dependent manner in vivo (Figure 2A), both proteins were purified via their N-terminal His_6_-tags (Figure S2) and subjected to *in vitro* analysis of their conformational dynamics using size-exclusion chromatography (SEC) and intrinsic tryptophan fluorescence spectroscopy at pH 7.4 and pH 5.8. SEC analysis revealed that both purified proteins show a distinctive peak corresponding to the size of a monomer, but at pH 5.8, wild-type AdiY exhibited a shift of approximately 0.3 mL toward lower retention volumes compared to pH 7.4 (Figure 3A), suggesting an increase in hydrodynamic radius and/or changes in interdomain interactions. Calibration with molecular mass standards indicated that AdiY adopts a more compact conformation at pH 7.4. In contrast, the AdiY-H34A/H60A variant showed only minor shifts between the two pH conditions, suggesting a reduced sensitivity to pH-induced conformational rearrangements. Intrinsic tryptophan fluorescence measurements (excitation at 280 nm, emission recorded between 300 and 400 nm) supported these findings (Figure 3B-D). It is noteworthy that the fluorophores are distributed unevenly across the domains: the N-terminal region contains two Trp and two Tyr, whereas the C-terminal region contains one Trp and seven Tyr. This enables domain-aware interpretation of the intrinsic fluorescence signals.

**Figure 3:**
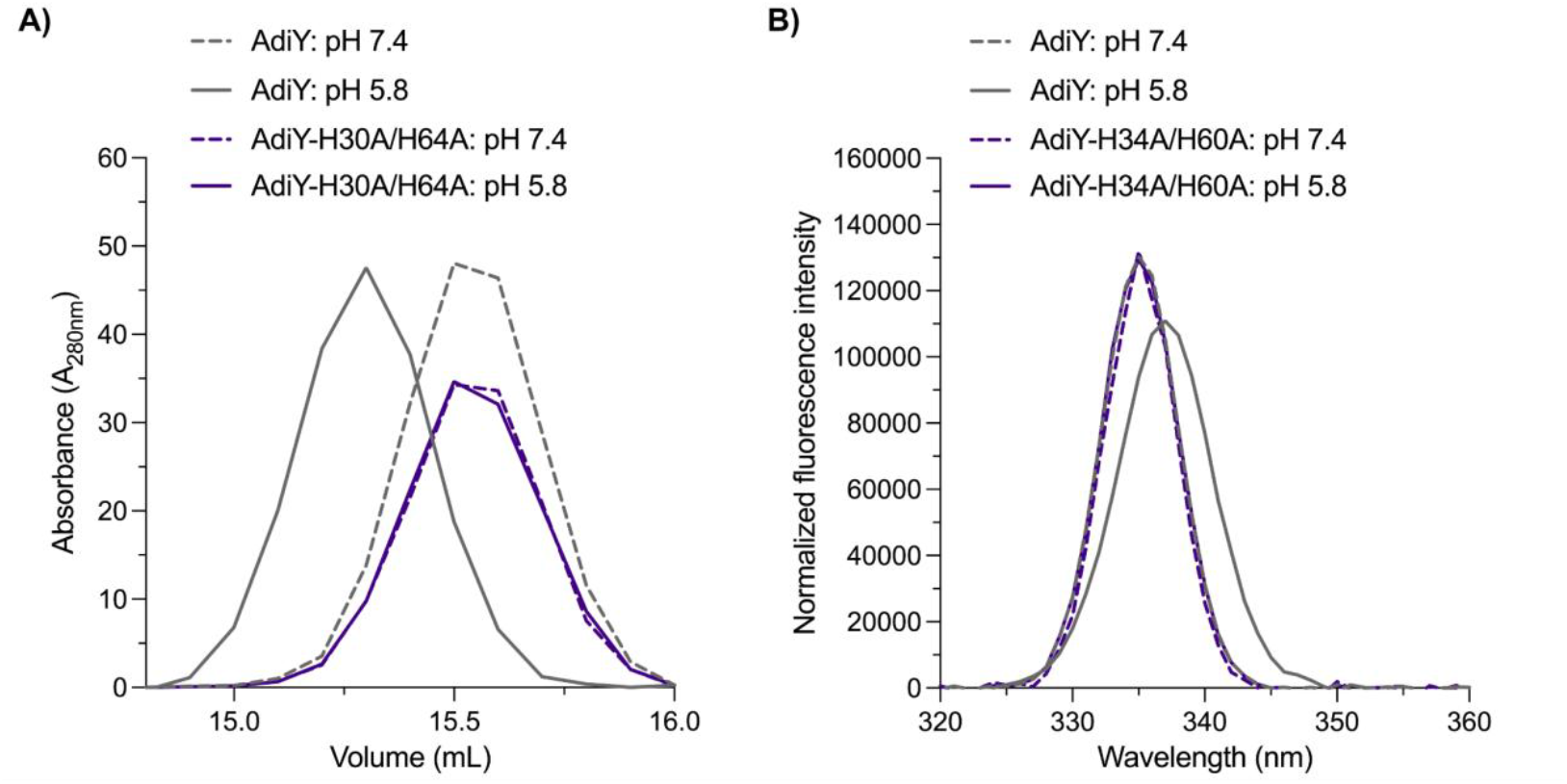
Comparative analysis of AdiY at pH 7.4 and pH 5.8 by size exclusion chromatography (SEC) and intrinsic tryptophan fluorescence spectroscopy. (**A**) SEC elution profiles of wild-type AdiY and the AdiY-H34A/H60A variant at pH 7.4 and pH 5.8. **(B)** Intrinsic tryptophan fluorescence emission spectra (excitation at 280 nm) of wild-type AdiY and the AdiY-H34A/H60A variant at pH 7.4 and pH 5.8.

At pH 7.4, wild-type AdiY showed an emission maximum at approximately 335–336 nm. At lower pH (5.8) we observed a red shift (∼2 nm) of the emission maximum and a decrease in fluorescence intensity, suggesting increased solvent exposure of tryptophan residues due to pH-induced structural rearrangements (Figure 3B). The AdiY-H34A/H60A variant showed minimal spectral shifts and intensity changes compared to wild-type AdiY at pH 7.4 (Figure 3C). At pH 5.8, its emission maximum remained at approximately 335–336 nm (Figure 3D), similar to that of wild-type AdiY at pH 7.4, further supporting a reduced conformational flexibility in the mutant relative to the wild-type protein.

Together, these data indicate that AdiY experiences significant pH-dependent conformational transitions (Figure 2B), that do not occur in the AdiY-H34A/H60A variant.

### AdiY-DNA interaction is pH-dependent in vitro

Given the pH-dependent conformational changes observed in AdiY wild-type (Figure 3), we investigated whether these structural shifts influence the ability of AdiY wild-type to bind the target promoters *in vitro*, in comparison to the AdiY-H34A/H60A variant. We used purified AdiY wild-type and AdiY-H34A/H60A proteins to quantitatively determine their binding affinity to the *adiA, adiC*, and *adiY* promoters using surface plasmon resonance (SPR) spectroscopy at different pH values (Figure 4). At pH 6.0, wild-type AdiY binds preferentially to the *adiA* and *adiC* promoters, with affinities of 170 ± 23.2 nM and 120 ± 34.4 nM, respectively (Figure 4A and Table 1). In both cases, the stoichiometry was 1:4 (DNA:protein), indicating that AdiY assembles as a tetramer on the promoter DNA (Figure 4A and Table 1). Furthermore, AdiY bound the *adiA* and *adiC* promoters with similar association rate constants (*k*_a_), averaging 1.01 × 10^4^ M^−1^ s^−1^ and 1.60 × 10^4^ M^−1^ s^−1^, respectively. The dissociation rates (*k*_d_) were also comparable at 1.70 × 10^3^ s^−1^ for the *adiA* and 1.80 × 10^3^ s^−1^ for the *adiC* promoter, indicating that once bound, the complexes had similar stability (Table 1). No binding of wild-type AdiY to the *adiY* promoter was detected under the same conditions or at any tested pH values (Figure 4A and S3). The AdiY-H34A/H60A mutant failed to exhibit any detectable DNA binding across all tested conditions (Figure 4B and S4), underscoring the critical role of histidine residues at positions 34 and 60 in mediating the pH-dependent conformational changes required for promoter binding.

**Table 1:** Kinetic parameters of AdiY binding to its target promoter determined by surface plasmon resonance (SPR). Association rate (*k*_a_) and dissociation (*k*_d_) rate constants and the equilibrium dissociation constants (K_D_) were determined for AdiY binding to the *adiA* and *adiC* promoter fragments at the indicated pH values. Values represent means from one or more independent experiments (N), with standard deviations (SD) where applicable. Measurements were performed using increasing concentrations of His_6_-AdiY (31.25 nM to 2000 nM) immobilized on a DNA-coated sensor chip. Binding to the *adiY* promoter was not detectable and is therefore not included in the table. Stoichiometry refers to DNA:protein ratios.

| Promoter | pH | Association constant [M<sup>-1</sup> s<sup>-1</sup>] |  |  | Dissociation constant [s<sup>-1</sup>] |  |  | Equilibrium constant [M] |  |  | Stoichiometry |
| --- | --- | --- | --- | --- | --- | --- | --- | --- | --- | --- | --- |
|  |  | Mean | SD | N | Mean | SD | N | Mean | SD | N |  |
| *adiA* | 5.8 | 1.52×10<sup>4</sup> |  | 1 | 2.11×10<sup>3</sup> |  | 1 | 1.39×10<sup>-7</sup> |  | 1 | 1:4 |
| *adiA* | 6.0 | 1.01×10<sup>4</sup> | 1.53×10<sup>3</sup> | 3 | 1.70×10<sup>3</sup> | 1.15×10<sup>4</sup> | 3 | 1.70×10<sup>-7</sup> | 2.32×10<sup>-8</sup> | 3 | 1:4 |
| *adiA* | 6.3 | 8.17×10<sup>3</sup> |  | 1 | 1.81×10<sup>3</sup> |  | 1 | 2.21×10<sup>-7</sup> |  | 1 | 1:3 |
| *adiA* | 6.5 | 1.62×10<sup>4</sup> |  | 1 | 4.59×10<sup>3</sup> |  | 1 | 2.83×10<sup>-7</sup> |  | 1 | 1:2 |
| *adiC* | 5.8 | 2.15×10<sup>4</sup> |  | 1 | 2.64×10<sup>3</sup> |  | 1 | 1.23×10<sup>-7</sup> |  | 1 | 1:4 |
| *adiC* | 6.0 | 1.60×10<sup>4</sup> | 5.72×10<sup>3</sup> | 3 | 1.80×10<sup>3</sup> | 2.46×10<sup>4</sup> | 3 | 1.20×10<sup>-7</sup> | 3.44×10<sup>-8</sup> | 3 | 1:4 |
| *adiC* | 6.3 | 1.29×10<sup>4</sup> |  | 1 | 1.53×10<sup>3</sup> |  | 1 | 1.18×10<sup>-7</sup> |  | 1 | 1:3 |
| *adiC* | 6.5 | 2.36×10<sup>3</sup> |  | 1 | 2.03×10<sup>2</sup> |  | 1 | 85.8×10<sup>-7</sup> |  | 1 | 1:2 |

**Figure 4.**
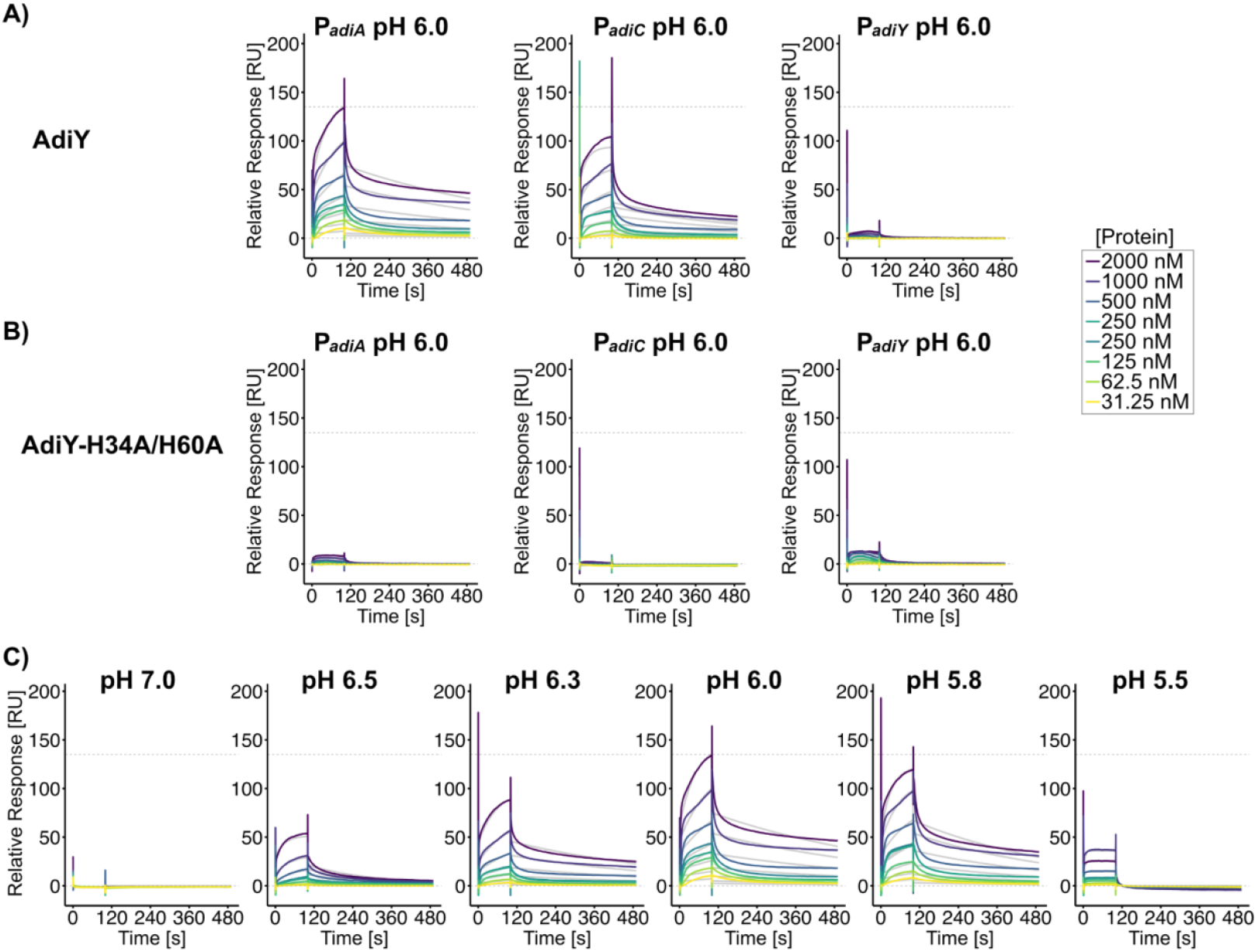
pH-dependent binding of AdiY and AdiY-H34A/H60A variant with target promoters analyzed by surface plasmon resonance (SPR) spectroscopy. **(A, B, C)** Biotinylated DNA fragments comprising the promoters of *adiA* (P_*adiA*_), *adiC* (P_*adiC*_), or *adiY* (P_*adiY*_) were captured on SA sensor chips. Solutions of purified AdiY wild-type **(A)** or purified AdiY-H34A/H60A **(B)** at pH 6.0 were passed over the sensor chip. **(C)** Solutions of purified AdiY wild-type were passed over the *adiA* promoter, immobilized on the sensor chip, at different pH values ranging from 7.0 to 5.5. Sensorgram color coding corresponds to increasing protein concentrations, as indicated in the panel on the right. The dashed line at 0 RU represents the baseline, while the dashed line at 135 RU marks the maximal response of wild-type AdiY to the *adiA* promoter at pH 6.0. Where applicable, association rate (*k*_a_) and dissociation rate (*k*_d_) constants and the equilibrium dissociation constants (K_D_) are summarized in Table 1.

To assess pH dependence *in vitro*, the binding of AdiY wild-type to its target promoter was tested across a pH range from 7.0 to 5.5. Binding of AdiY wild-type to the *adiA* and *adiC* promoters was strongest at acidic pH values (6.0, 5.8, and 6.3), weaker at pH 6.5 and undetectable at pH 5.5 or physiological pH values above 6.7. However, tetramerization of AdiY was only achieved at pH values of 5.8 and 6.0 on these two promoters (Figure 4C, Table 1, and S3). Across the tested pH range, *k*_a_ values for the *adiA* promoter varied from 8.17 × 10^3^ M^−1^ s^−1^ at pH 6.3 to 1.62 × 10^4^ M^−1^ s^−1^ at pH 6.5, whereas the value for the *adiC* promoter showed a slightly higher maximum *k*_a_ (2.15 × 10^4^ M^−1^ s^−1^ at pH 5.8) which was lower at pH 6.5 (*k*_a_ 2.36 × 10^3^ M^−1^ s^−1^) (Table 1). For both promoters, the dissociation rate was lowest and binding stability was highest at mildly acidic pH values (5.8–6.3). At pH 5.8, the *adiA* and *adiC* promoters exhibited *k*_d_ values of 2.11 × 10^3^ s^−1^ and 2.64 × 10^3^ s^−1^, respectively. At pH 6.5, the dissociation rates increased for the *adiA* promoter (4.59 × 10^3^ s^−1^) and decreased markedly for the *adiC* promoter (2.03 × 10^2^ s^−1^) (Table 1). Overall, both promoters exhibited the fastest association and slowest dissociation at moderately acidic pH values, with maximal binding between pH 5.8 and 6.3.

These results demonstrate that AdiY binds specifically and cooperatively to the *adiA* and *adiC* promoters, with DNA-binding activity sharply tuned to a narrow range of moderately acidic intracellular pH values. Outside this range, binding was almost completely abolished, indicating that AdiY is selectively activated under these conditions.

## Discussion

This study identified AdiY as a direct pH-sensor and transcriptional activator of *adiA* and *adiC* of the AR3/Adi system in *E. coli*. AdiY is activated by protonation of two histidine residues at positions 34 and 60 in the N-terminal domain, which triggers a conformational change of the position of the C-terminal domain, leading to DNA-binding and tetramerization. Substitution of His34 and His60 not only abolished the pH sensitivity but also the pH-dependent conformational rearrangement of AdiY. SPR analyses demonstrated that wild-type AdiY binds specifically and cooperatively to the *adiA* and *adiC* promoters *in vitro*, with maximum affinity and tetramerization observed at mildly acidic pH values between pH 5.8 and 6.3 (Figure 4, Figure S3, Table 1). Outside this pH range, AdiY did not bind to the DNA, suggesting its very restricted activation. This observed pH dependence is consistent with the activation profile of the Adi system, which reaches its maximum at a low external pH between 5.0 and 4.4, a condition that causes a decrease of the cytoplasmic pH to about 5.8 to 6.0 (27). Under these conditions, AdiY senses the intracellular acidification, tetramerizes and binds to its target promoters, and induces expression of *adiA* and *adiC*.

Based on these findings, we propose the following model (Figure 5): (i) At neutral pH, AdiY predominantly exists as an inactive monomer. (ii) Acidification of the cytoplasm induces protonation of key histidines, which triggers a conformational change. (iii) Following promoter recognition, AdiY undergoes tetramerization, which stabilizes the transcription complex. (iv) This structural rearrangement enables pH-dependent activation of the target genes *adiA* and *adiC* of the Adi system.

**Figure 5.**
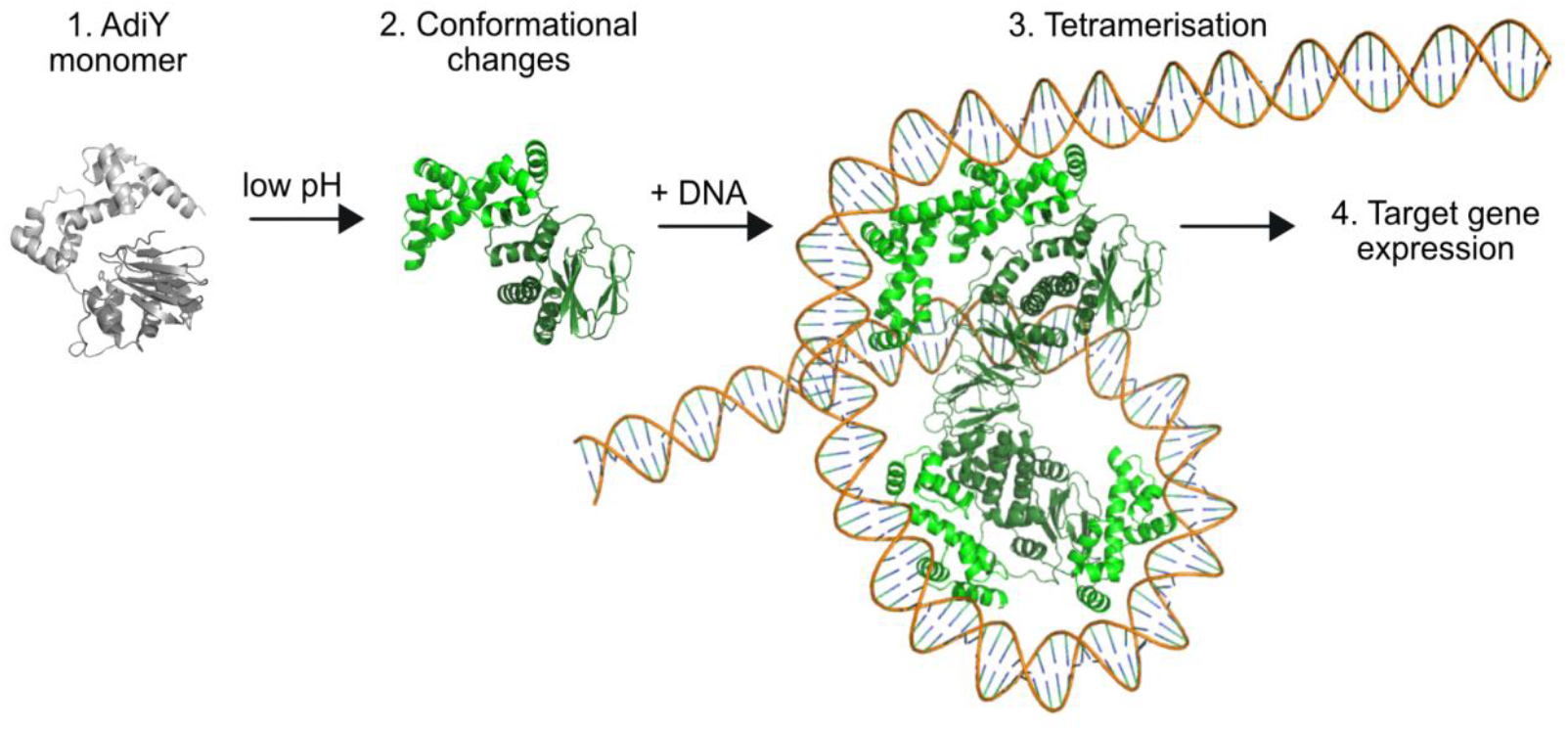
pH-dependent activation mechanism of AdiY. Schematic model illustrating the proposed activation pathway of the AdiY transcription factor in *Escherichia coli* under acidic stress. (1) Under neutral pH, AdiY exists predominantly as an inactive monomer (grey). (2) Low cytoplasmic pH induces protonation of key histidine residues, which triggers a conformational change. (3) Upon binding to DNA, AdiY undergoes tetramerization, which stabilizes the transcription complex. (4) This structural rearrangement enables pH-dependent activation of the target genes *adiA* and *adiC* of the Adi system. Structural predictions were generated by Alphafold 3. The figure was prepared with PyMOL (v3.1.4.1).

The activity of the arginine decarboxylase AdiA is also strongly affected by the intracellular pH, which regulates its oligomerization. At pH 5.2, AdiA forms active decamers, which dissociate in inactive dimers at pH >6.0 (32). It is noteworthy that the catabolic genes for arginine are downregulated under severe acid stress (33), presumably to preserve arginine for decarboxylation. In summary, all components of the Adi system are finely tuned to be most active under severe acid stress within a very narrow range and become immediately inactive when the intracellular pH rises. The protonation of amino acids whose side chain pKa values are around pH 6.0 (histidine) or below (aspartic acid, pKa 3.88, glutamic acid, pKa 4.25) is a well-documented molecular mechanism in bacterial acid stress sensing (26, 34, 35), exemplified by the histidine kinases SsrA (five histidines), ArsS (one histidine, four glutamic acids, and the ToxR-like regulator CadC (three aspartic acids, two glutamic acids) (5, 6, 35–38). A difference between the pH sensors lies in the compartment they monitor. AdiY responds to cytoplasmic acidification, where the protonation of histidine residues at positions 34 and 60 causes a conformational change that enables DNA binding (Figure 5). In contrast, SsrA, AsrS, and CadC detect acidity in the periplasm, e.g., CadC via protonation of a cluster of negatively charged amino acids (Asp198, Asp200, Glu461, Glu468, Asp471) in its periplasmic domain, which is located at the interface between two monomers (26). Cytoplasmic regulators such as AphB of *Vibrio cholerae* (39) and *V. campbellii* (40) also integrate intracellular pH into the regulation of virulence and stress responses. However, unlike AdiY, *V. cholerae* AphB relies on cysteine- and lysine-mediated redox and protonation changes (39). *V. campbellii* AphB achieves regulation through rapid, high-affinity promoter binding (40), while AdiY relies on cooperative, tetramerization-driven DNA binding with sub-micromolar affinity, which is restricted to a narrow pH range. The requirement for tetramerization to achieve stable promoter occupancy is consistent with the mechanisms described for AraC and LacI family regulators, where oligomerization is essential for precise DNA binding and transcriptional control (41, 42).

AdiY belongs to the large family of AraC/XylS family of transcription regulators. Most of them consist of a characteristic and conserved DNA-binding domain and a “companion” domain (43), also referred to as an “effector” or “regulatory” domain (13). In many proteins, this domain has been shown to sense specific signaling molecules, such as fatty acids (VirF), arabinose (AraC), or xylose (XylS). When these molecules are bound, the transcription factors bind as a dimer to the corresponding promoter, inducing transcription (13). AdiY is an exception in this family, as it undergoes protonation/deprotonation and forms a tetramer (Figure 5).

The Adi system is also found in other enterobacteria, such as *Shigella, Salmonella*, and *Citrobacter* (6). *E. coli* AdiY is highly similar to its orthologs in *Escherichia albertii* and *Shigella*, all of which retain His34 and His60 (Figure S5). AdiY in representative strains of *Salmonella* and *Citrobacter* has His34 but no His60, where an arginine is located instead. It is worth noting that *Salmonella* completely lacks the Gad system, and *Citrobacter* lacks many regulatory components of the Gad system. In comparison, *E. coli* and *Shigella* have four acid-inducible amino acid–dependent decarboxylase systems and the highest resistance under extreme acid stress (6). Therefore, we hypothesize that the histidine-controlled protonation of AdiY in *E. coli* has evolved as a highly restrictive mechanism that responds to minute pH changes in the cytosol and confers maximal resistance for *E. coli*, for example, during passage through the strongly acidic environment of the stomach.

From an applied perspective, understanding this histidine-dependent pH-sensing mechanism opens up new avenues for developing AdiY in a pH-sensor that is useful in biosensors for microbiological, clinical and biotechnological applications.

## Materials and Methods

### Construction of plasmids and strains

Molecular methods were performed according to standard protocols or as instructed by the manufacturer. Kits for the isolation of plasmids and the purification of PCR products were purchased from Süd-Laborbedarf (SLG; Gauting, Germany). Enzymes and HiFi DNA Assembly Master Mix were purchased from New England BioLabs (Frankfurt, Germany). Strains, plasmids and primers used in this study are summarized in Tables 2 and 3.

**Table 2.**
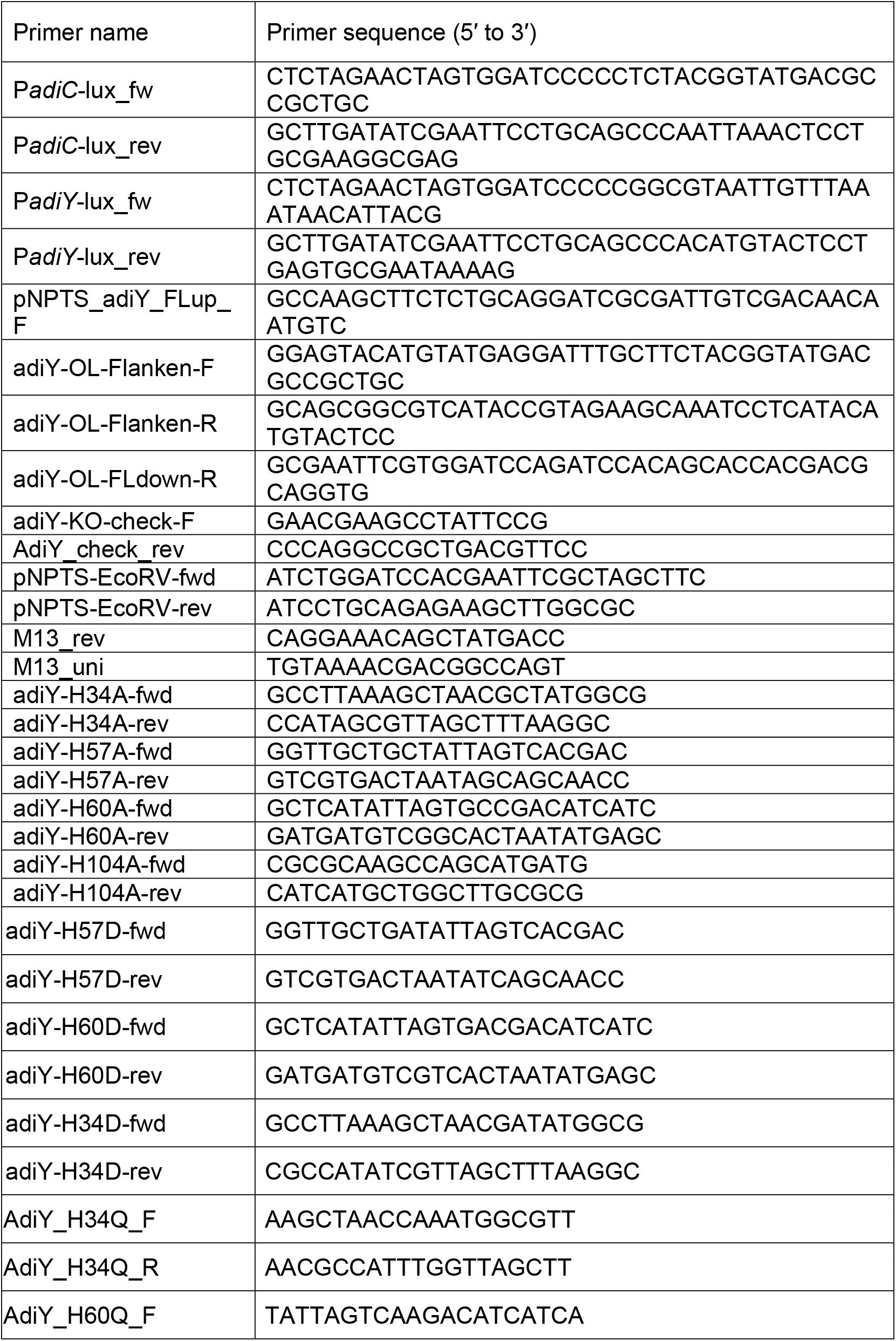
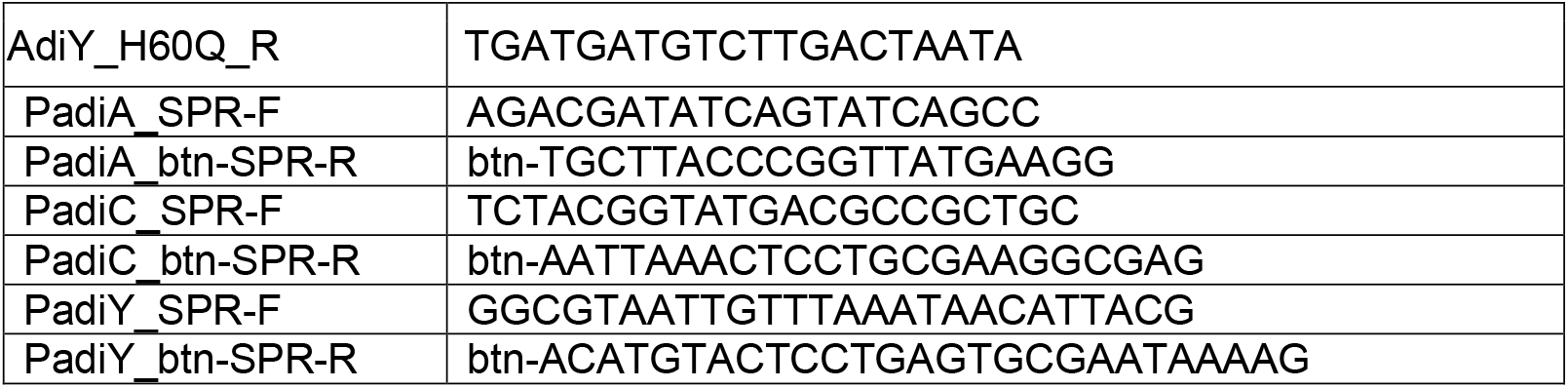
Primers sequences used in this study. Btn, biotin.

**Table 3:**
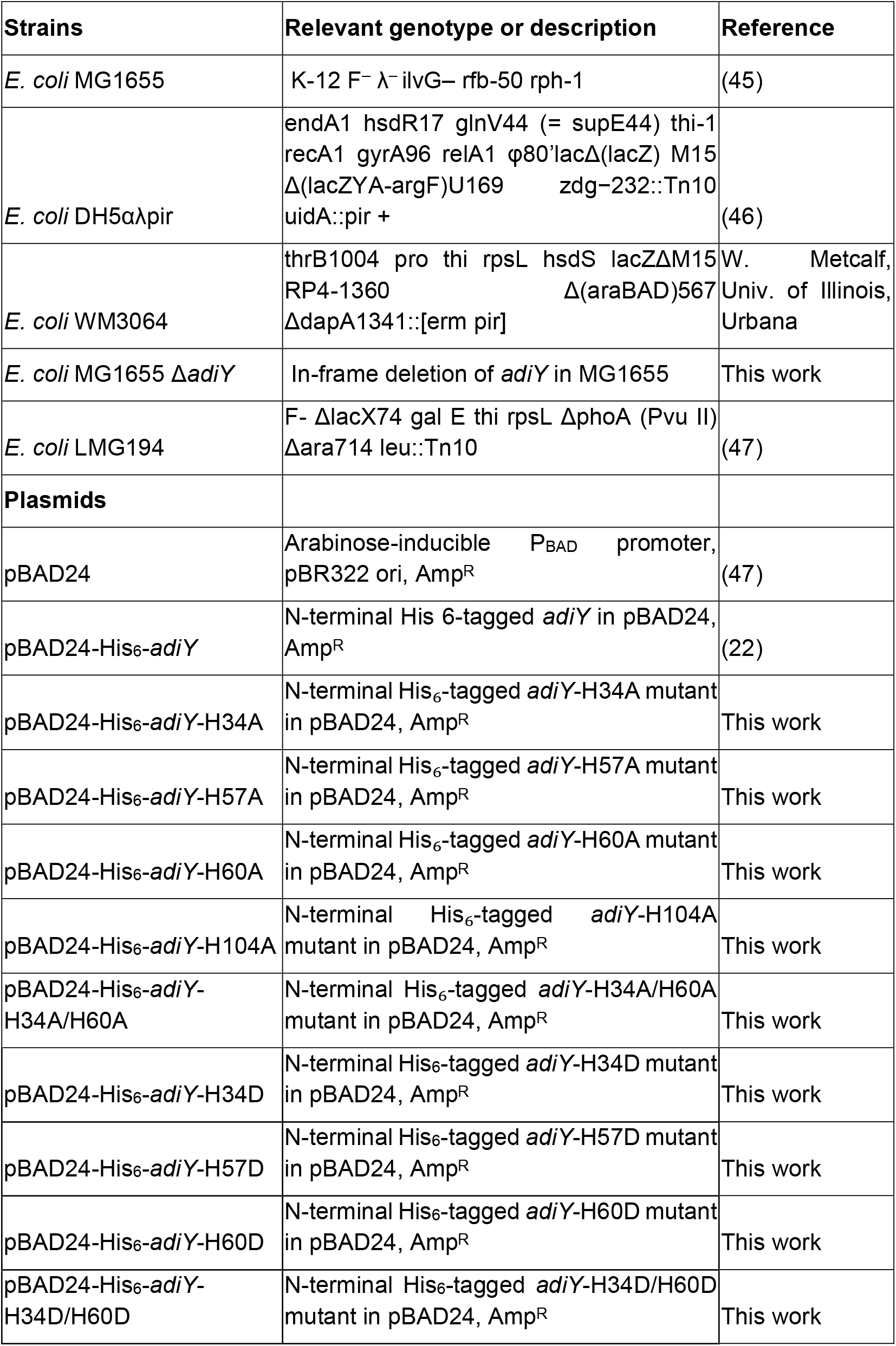
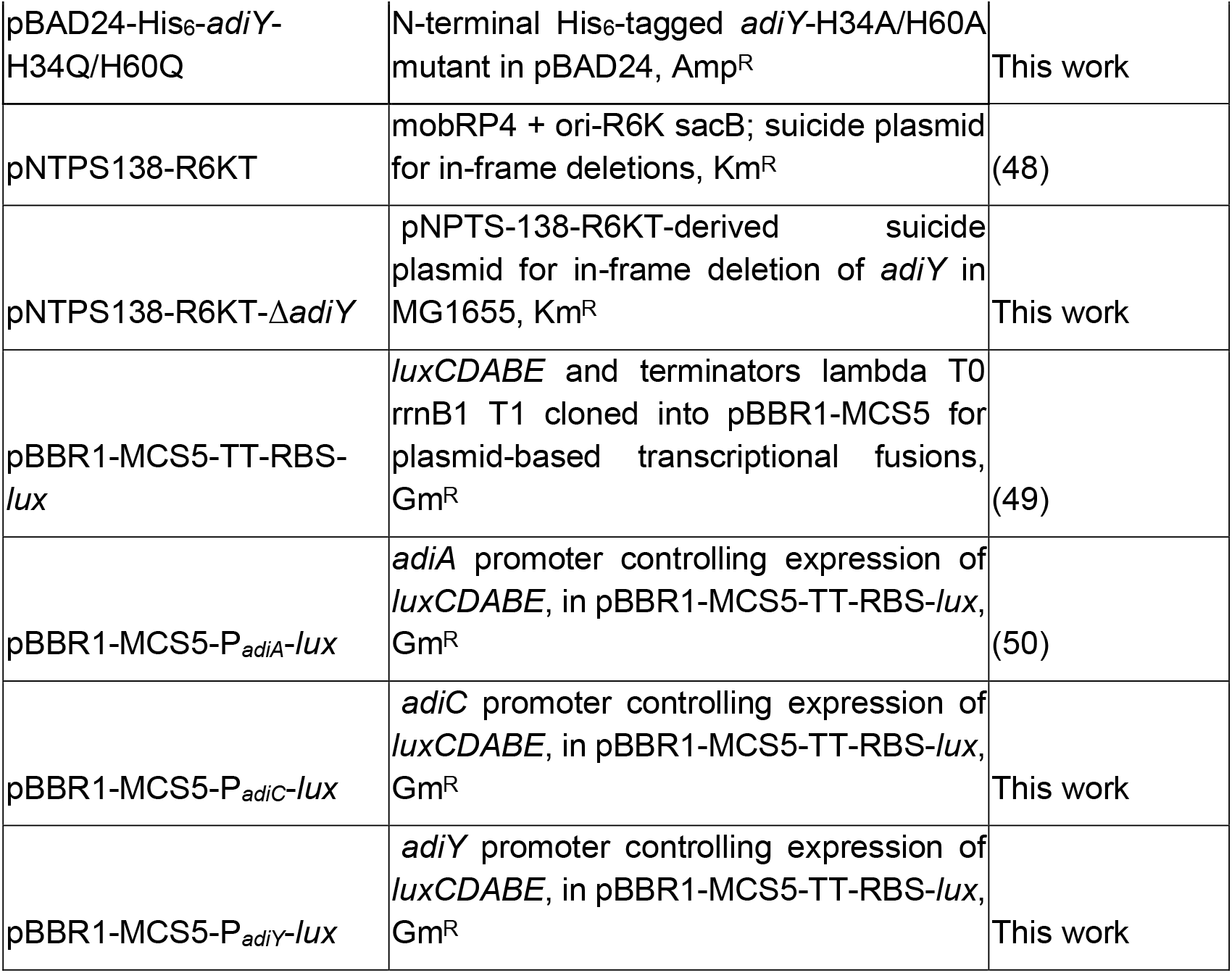
Strains and plasmids used in this study.

To construct the reporter plasmids pBBR1-MCS5-P_*adiC*_-*lux* and pBBR1-MCS5-P_*adiY*_-*lux*, 200 bp and 300 bp fragments from the regions upstream of *adiC* or *adiY*, respectively, were amplified by PCR using the appropriate primer pairs (PadiC-lux_fw and PadiC-lux_rev, or PadiY-lux_fw and PadiY-lux_rev). Genomic DNA from *E. coli* strain MG1655 served as the template. After purification, the PCR products were each assembled into SmaI-digested pBBR1-MCS5-TT-RBS-lux plasmid using Gibson assembly (44), resulting in the plasmids pBBR1-MCS5-P_*adiC*_-*lux* and pBBR1-MCS5-P_*adiY*_-*lux*. Colony PCR and sequencing verified the correct insertion.

To construct a marker-less in-frame deletion of *adiY* in *E. coli* MG1655, firstly, the suicide plasmid pNPTS138-R6KT-Δ*adiY* was generated as previously described in (22). Briefly, flanking regions of each 600 bp upstream and downstream of *adiY* were amplified by PCR from *E. coli* MG1655 genomic DNA using the appropriate primer pairs (pNPTS_adiY_FLup_F and adiY-OL-Flanken-R, or adiY-OL-Flanken-F and adiY-OL-FLdown-R). After purification, the PCR products were assembled into the PCR linearized pNPTS138-R6KT plasmid, previously amplified by PCR using the primer pair pNPTS-EcoRV-fwd and pNPTS-EcoRV-rev, via Gibson assembly (44). The resulting plasmid, pNPTS138-R6KT-Δ*adiY*, was verified by sequence analysis using primers M13_rev and M13_uni. To generate the deletion of *adiY* in *E. coli* MG1655, at first, the suicide plasmid pNPTS138-R6KT-Δ*adiY* was introduced into *E. coli* MG1655 by conjugative mating using *E. coli* WM3064 as a donor in LB medium containing DAP. Single-crossover integration mutants were selected on LB plates containing kanamycin but lacking DAP. Single colonies were grown for a day without antibiotics and plated onto LB plates containing 10% (w/v) sucrose but lacking NaCl to select for plasmid excision. Kanamycin-sensitive colonies were checked for targeted deletion by colony PCR using primers bracketing the site of the insertion. Deletion of *adiY* was verified by colony PCR and sequencing using the primers adiY-KO-check-F and AdiY_check_rev, resulting in the strain *E. coli* MG1655 Δ*adiY*.

Site-directed mutagenesis of the *adiY* gene was performed by PCR amplification using primers containing the desired mutations within their overhangs. The plasmid pBAD24-His_6_-*adiY* served as the template (22). A list of the mismatch-containing oligonucleotide primers is provided in Table 2. The resulting PCR products, encoding the intended amino acid substitutions, were assembled into SmaI-digested pBAD24 using Gibson assembly. Successful construction of the mutant plasmids was confirmed by colony PCR and Sanger sequencing.

### Structural predictions

Protein structure predictions were performed using AlphaFold 3 (51–53). The complete amino acid sequence was submitted to the AlphaFold 3 web server using default parameters. AlphaFold 3 applies a transformer-based architecture to capture residue–residue relationships and employs iterative diffusion-based refinement to generate convergent three-dimensional structures. Model outputs include atomic coordinates for all non-hydrogen atoms and confidence metrics such as the predicted Local Distance Difference Test (pLDDT) scores and Predicted Aligned Error (PAE) maps. The model with the highest overall confidence was selected for further analysis. For visualization, the final models were loaded into PyMOL (v3.1.4.1) to inspect domain organization, substructure elements, and potential interaction interfaces. Model reliability was assessed using per-residue pLDDT scores, color-coded to indicate prediction confidence (blue: very high, cyan: high, yellow: low, orange: very low). The AdiY-H34A/H60A model displayed high to very high confidence across most secondary structure elements, with reduced reliability restricted to flexible loop regions (see Figure S7). AlphaFold 3 was also applied in multimeric mode to predict AdiY– DNA interactions using the AdiY sequence and the adiA promoter fragment.

### *In vivo* AdiY activity assay

To evaluate the pH-dependent activity of AdiY and its histidine variants *in vivo*, we employed a bioluminescent reporter system based on a plasmid carrying the *adiA* promoter fused upstream of the *luxCDABE* operon. The reporter construct was transformed into *E. coli* MG1655 cells, which were subsequently co-transformed with expression plasmids encoding either wild-type AdiY or single histidine-to-alanine variants under the control of an arabinose-inducible promoter (pBAD24-derived vector backbone). Strains were grown overnight at 37°C in LB medium supplemented with the appropriate antibiotics, pH 7.0. The next day, the cells were adjusted to an OD_600_ of 0.05 in fresh Citrate buffer adjusted to pH 4.4 or pH 7.0 and grown until an OD_600_ of approximately 0.5 was reached. They were then incubated in 96-well plates under either aerobic (shaking at 300 rpm) or microaerobic (no shaking) conditions. Induction of the production of AdiY or its variants was achieved by adding L-(+)-arabinose to a final concentration of 0.1% (w/v) at the time of inoculation into the 96-well plates. Bioluminescence was recorded at 10-minute intervals over 20 hours using a Tecan Infinite F500 plate reader (Tecan, Crailsheim, Germany), and promoter activity was quantified as the maximum relative light units (RLU) measured after 2h.

To examine how external pH influences AdiY-dependent promoter activation, *E. coli* MG1655 wild-type and an *adiY* deletion mutant were each transformed with the reporter plasmids pBBR1-MCS5-P*adiA-lux*, pBBR1-MCS5-P*adiC-lux*, or pBBR1-MCS5-P*adiY-lux*. Cultures were grown in LB medium adjusted to pH values ranging from 4.0 to 7.0. Luminescence and growth (OD_600_) were measured every 10 minutes using a Tecan Infinite F500 plate reader (Tecan, Crailsheim, Germany). Data was calculated as relative light units (RLU), calculated as counts per second per milliliter per OD_600_. Data analysis was conducted using GraphPad Prism (version 10.4.1).

### AdiY production and purification

N-terminally His6-tagged wild-type AdiY and the AdiY-H34A/H60A variant were expressed in *E. coli* LMG194 transformed with the respective pBAD24-His_6_-*adiY* or pBAD24-His_6_-*adiY*-H34A/H60A plasmids. Cultures were grown in 1 L of terrific broth (TB; 20 g Bacto Tryptone, 24 g Bacto Yeast Extract, 4.65 mL 86% v/v glycerol, and 100 mL potassium phosphate buffer pH 7.4 per liter) at 37 °C to an OD_600_ of 0.6, then induced with 0.2% arabinose for 6 hours at 37 °C. Cells were harvested by centrifugation and pellets were stored at –80 °C until use. Cell pellets were resuspended in 100 mL of lysis buffer (300 mM NaCl, 50 mM sodium phosphate pH 7.2, 2 mM DTT, 0.2 mM PMSF, DNase I) and lysed using a high-pressure homogenizer. The lysate was clarified by sequential centrifugation (5,000 × g for 30 min at 4 °C to remove debris, followed by 70000 RCF for 1 h at 4 °C to collect the soluble fraction). Proteins were purified by Ni-IDA affinity chromatography (Protino, MACHEREY-NAGEL): after binding, the resin was washed with 10 column volumes of wash buffer (same as lysis buffer with 30 mM imidazole), and proteins were eluted in 1.5 column volumes of elution buffer (50 mM sodium phosphate pH 5.8, 300 mM NaCl, 2% glycerol, 2 mM DTT, 5 mM EDTA). Fractions containing AdiY were pooled and further purified by size-exclusion chromatography (SEC) using a HiLoad 16/600 Superdex 75 pg column (Cytiva) equilibrated in 50 mM sodium phosphate pH 5.8 (or 7.2), 300 mM NaCl, 2% (v/v) glycerol, 2 mM DTT, and 2 mM EDTA. The column was calibrated with molecular weight standards (Aprotinin, Ribonuclease A, Carbonic anhydrase, Ovalbumin, Conalbumin; Cytiva). Only the fractions corresponding to the AdiY monomer, as determined by comparison to the calibration curve, were collected for subsequent analysis. Final protein samples were concentrated using Vivaspin^®^ 20 (10 kDa cutoff, Merck), quantified, aliquoted, and stored at –20°C. Protein expression and purification were monitored by SDS–PAGE (12% polyacrylamide gels; 10 µg total protein per lane; PageRuler Prestained Protein Ladder, Thermo Scientific) and Western blotting. Proteins were transferred onto PVDF membranes (0.45 µm, Millipore) and probed with anti-His monoclonal antibodies (1:5,000 dilution; Sigma-Aldrich), followed by HRP-conjugated secondary antibodies (1:10,000 dilution). Signals were visualized using enhanced chemiluminescence (ECL, Cytiva). Representative results are shown in Supplementary Figure S2.

### Conformational analysis of AdiY by size exclusion chromatography (SEC)

The pH-dependent conformational changes of AdiY were analyzed by size-exclusion chromatography (SEC) using a Superdex 75 Increase 10/300 GL column (Cytiva), pre-equilibrated with 50 mM sodium phosphate buffer (pH 5.8 or 7.4) containing 300 mM NaCl, 2% (v/v) glycerol, 2 mM DTT, and 2 mM EDTA. SEC was performed on an ÄKTA Pure system with a flow rate of 0.5 mL/min, using a 0.5 mL protein sample at a concentration of 0.5 mg/mL. The column was calibrated with molecular weight standards as described above. Elution profiles were analyzed using UNICORN 7.0 software, and the apparent molecular weight of AdiY was determined by comparing its elution volume to the standard calibration curve.

### Conformational analysis of AdiY by intrinsic tryptophan fluorescence spectroscopy

Intrinsic tryptophan fluorescence measurements were conducted using a FluoroMax-3 (Horiba) at 25 °C. The excitation wavelength was set to 280 nm to excite tryptophan residues selectively, and emission spectra were recorded from 300 to 400 nm. Spectra were collected at 1 nm intervals with an integration time of 1 s per increment and slit widths of 5 nm for both excitation and emission. Purified wild-type AdiY and the AdiY--H34A/H60A were prepared at a final concentration of 1 μM in 50 mM sodium phosphate buffer at pH 5.8 or 7.4, supplemented with 300 mM NaCl, 2% (v/v) glycerol, and 2 mM DTT. Fluorescence spectra were recorded for each protein under both buffer conditions to assess pH-dependent structural rearrangements. For comparative analysis, bovine serum albumin (BSA, 5 μM; Sigma-Aldrich) was used as a control under identical experimental conditions. To reduce inner filter effects, fluorescence intensities were corrected for absorption of the exciting light and reabsorption of the emitted light using the equation F_corr_=F_obs_×10^(Aexc+Aem)/2^, where F_corr_ is the corrected fluorescence, F_obs_ the measured fluorescence, A_exc_ the absorbance at the excitation wavelength, and A_em_ the absorbance at each emission wavelength; absorbance values were obtained from UV-Vis spectra of the same samples measured under identical conditions(36, 54). Fluorescence data were analyzed in GraphPad Prism 10.4.1 to quantify spectral shifts and changes in intensity. All experiments were performed in triplicate to ensure reproducibility.

### Surface plasmon resonance spectroscopy

Surface plasmon resonance (SPR) experiments were performed on a Biacore T200 instrument at 25°C using a carboxymethyl dextran sensor chip precoated with streptavidin (SA Sensor Chip Series S; Cytiva). All experiments were performed at 25 °C using a running buffer composed of 50 mM sodium phosphate (pH 6.0–7.0), 250 mM NaCl, 2% (v/v) glycerol, 2 mM DTT, and 0.005% (v/v) detergent P20. For measurements at pH 5.5 and pH 5.8, sodium phosphate was replaced with 50 mM citrate, while all other buffer components and conditions were kept identical. pH 6.0 was used as the standard condition and was included in all experiments to facilitate direct comparison between SPR runs. To immobilize DNA fragments comprising the promoters of *adiA, adiC*, and *adiY* (each 200 bp), these promoter regions were amplified by PCR using *E. coli* MG1655 genomic DNA as the template with the following primers to incorporate a biotin (btn) tag PadiA_SPR-F and PadiA_btn-SPR-R, PadiC_SPR-F and PadiC_btn-SPR-R, PadiY_SPR-F and PadiY_btn-SPR-R, respectively. Before immobilizing the DNA fragments comprising the btn-tagged promoters of *adiA, adiC*, and *adiY*, the chip was equilibrated by three injections of 1 M NaCl/50 mM NaOH applied at a flow rate of 10 mL/min. Next, the biotinylated promoter fragments (10 nM) were injected at a flow rate of 10 mL/min for a total contact time of 240 s on flow cells 2, 3, and 4, respectively. The chips were then washed by injecting 1 M NaCl, 50 mM NaOH, and 50% (v/v) propan-2-ol. About 150 RU (response units) of the promoter fragments were bound per flow cell. Kinetic analyses of the interactions between purified wild-type His_6_-AdiY and the His_6_-AdiY-H34A/H60A variant with promoter fragments were conducted at 25 °C in running buffer at various pH values, using a flow rate of 30 mL/min. Seven concentrations of either wild-type AdiY or the AdiY-H34A/H60A variant (two-fold serial dilutions from 2000 nM down to 31.25 nM with a repetition of 62.5nM), dissolved in each running buffer, were passed over the flow cells for 120 s, and the complexes formed were allowed to dissociate for 360 s before the next cycle started. After each cycle, the surface was regenerated by injection of 2.5 M NaCl for 30 s, followed by 0.5% (wt/v) SDS for 30 s, at a flow rate of 30 mL/min. Sensorgrams were recorded using Biacore T200 Control software version 2.0.2 and analyzed with Biacore T200 Evaluation software version 3.2.2. The surface of flow cell 1 was not coated and was used to obtain blank sensorgrams for subtraction of the bulk refractive index background. The referenced sensorgrams were normalized to a baseline of 0. Peaks in the sensorgrams at the beginning and end of the injection are due to the run-time difference between the flow cells for each chip. To calculate the association and dissociation rate constants as well as the equilibrium dissociation constants (K_D_), the sensorgram curves were used and the kinetics were fitted assuming a Langmuir 1:1 binding model using Biacore T200 Evaluation software version 3.2.2. Visualisation of the sensograms was performed in R (R Core Team, 2024 (55)) using the tidyverse and ggplot2 (56) packages within the RStudio IDE (57).

### Identification and alignment of AdiY homologs

AdiY homologs were identified with BLASTp (58) against the NCBI RefSeq Select (59) protein database (downloaded September 2025), using the *E. coli* K-12 MG1655 AdiY sequence as the query. Hits were retained if they met an e-value ≤ 1×10^−^100^−100^ and were within ±10% of the query length. Partial sequences were excluded. To avoid redundancy, we maintained a single representative per species, resulting in 17 non-redundant orthologs. Sequences were aligned in CLC Main Workbench v24.0.1 (QIAGEN) with the progressive “High accuracy” setting (gap-open cost = 10; gap-extension cost = 1; default protein matrix), treating terminal gaps as internal (60). No manual editing was performed. Accession numbers and organism names are provided in Table S1, and residue positions in the figure are numbered relative to *E. coli* AdiY.

## Acknowledgments

We thank Sabine Peschek for excellent technical assistance and Leen Houri for the construction of plasmid-encoded AdiY variants carrying alanine substitutions at positions 34 and 60.

